# Classification and Specific Primer Design for Accurate Detection of SARS-CoV-2 Using Deep Learning

**DOI:** 10.1101/2020.03.13.990242

**Authors:** Alejandro Lopez-Rincon, Alberto Tonda, Lucero Mendoza-Maldonado, Daphne G.J.C. Mulders, Richard Molenkamp, Carmina A. Perez-Romero, Eric Claassen, Johan Garssen, Aletta D. Kraneveld

## Abstract

In this paper, deep learning is coupled with explainable artificial intelligence techniques for the discovery of representative genomic sequences in SARS-CoV-2. A convolutional neural network classifier is first trained on 553 sequences from available repositories, separating the genome of different virus strains from the Coronavirus family with considerable accuracy. The network’s behavior is then analyzed, to discover sequences used by the model to identify SARS-CoV-2, ultimately uncovering sequences exclusive to it. The discovered sequences are first validated on samples from other repositories, and proven able to separate SARS-CoV-2 from different virus strains with near-perfect accuracy. Next, one of the sequences is selected to generate a primer set, and tested against other state-of-the-art primer sets on existing datasets, obtaining competitive results. Finally, the primer is synthesized and tested on patient samples (n=6 previously tested positive), delivering a sensibility similar to routine diagnostic methods, and 100% specificity. In this paper, deep learning is coupled with explainable artificial intelligence techniques for the discovery of representative genomic sequences in SARS-CoV-2. A convolutional neural network classifier is first trained on 553 sequences from NGDC, separating the genome of different virus strains from the Coronavirus family with accuracy 98.73%. The network’s behavior is then analyzed, to discover sequences used by the model to identify SARS-CoV-2, ultimately uncovering sequences exclusive to it. The discovered sequences are validated on samples from NCBI and GISAID, and proven able to separate SARS-CoV-2 from different virus strains with near-perfect accuracy. Next, one of the sequences is selected to generate a primer set, and tested against other state-of-the-art primer sets, obtaining competitive results. Finally, the primer is synthesized and tested on patient samples (n=6 previously tested positive), delivering a sensibility similar to routine diagnostic methods, and 100% specificity. The proposed methodology has a substantial added value over existing methods, as it is able to both identify promising primer sets for a virus from a limited amount of data, and deliver effective results in a minimal amount of time. Considering the possibility of future pandemics, these characteristics are invaluable to promptly create specific detection methods for diagnostics.

## Introduction

The Coronaviridae family presents a positive sense, single-strand RNA genome. These viruses have been identified in avian and mammal hosts, including humans. Coronaviruses have genomes from 26.4 kilo base-pairs (kbps) to 31.7 kbps, with G + C contents varying from 32% to 43%; human-infecting coronaviruses belonging to this family include SARS-CoV, MERS-CoV, HCoV-OC43, HCoV-229E, HCoV-NL63 and HCoV-HKU1^1^. In December 2019, SARS-CoV-2, a novel, human-infecting Coronavirus was identified in Wuhan, China, using Next Generation Sequencing (NGS)^2^. As of the 12^*th*^ August of 2020, the new SARS-CoV-2 has 20,162,474 confirmed cases across almost all countries, with 3,641,603 cases in the European region^3^. In addition, SARS-CoV-2 has an estimated mortality rate of 3-4%, and it is spreading faster than SARS-CoV and MERS-CoV^4^.

As a typical RNA virus, new mutations appears every replication cycle of Coronavirus, and its average evolutionary rate is roughly 10-4 nucleotide substitutions per site each year^2^. In the specific case of SARS-CoV-2, RT-qPCR testing using primers in ORF1ab and N genes have been used to identified the infection in humans^5^. This method has come into question; Yang et al. in a study from 866 respiratory specimens showed that for 0-7 days after onset of illness, the sputum samples had a negative rate of 11.1% in severe and 17.8% in mild cases, follow by 26.7% and 27.0% in nasal swabs and finally 40% and 38.7% for throat swabs^6^. Zhao et al. reports that 35.2% of 173 patients did not show positive in RT-PCR test^7^, which has been further explored by Arevalo et al.^8^ and Woloshin et al.^9^. These problems could be the result of the variation of viral RNA sequences within virus species, and the viral load in different anatomic sites^10^. It has been noted that, population mutation frequency of site 8,872 located in ORF1ab gene and site 28,144 located in ORF8 gene gradually increased from 0 to 29% as the epidemic progressed^11^. Apart from the false negative test problems, SARS-CoV-2 assays can yield a small portion of false positives through nonspecific detection of other Coronaviruses, as the virus is closely related to other Coronavirus organisms^12^. In addition, SARS-CoV-2 may be present with other respiratory infections, hindering its identification^13,14^.

Thus, it is fundamental to improve existing diagnostic tools to contain the spread. For example, diagnostic tools combining computed tomography (CT) scans with deep learning have been proposed, achieving an improved detection accuracy of 82.9%^15^. Another solution being used for studying SARS-CoV-2, is sequencing of the viral complementary DNA (cDNA). For example, we can use this sequencing data with cDNA, resulting from the PCR of the original viral RNA; e,g, Real-Time PCR amplicons to identify the SARS-CoV-2^16^.

Classification using viral sequencing techniques is mainly based on alignment methods such as FASTA^17^ and BLAST^18^. These methods rely on the assumption that cDNA sequences share common features, and their order prevails among different sequences^19,20^. However, these methods suffer from the necessity of needing base sequences for the detection^21^. Nevertheless, it is necessary to develop innovative improved diagnostic tools that target the genome to improve the identification of pathogenic variants, as sometimes several tests, are needed to have an accurate diagnosis. Therefore, as an alternative, deep learning methods have been suggested for classification of DNA sequences. The advantage of these methods are that they do not need pre-selected features to identify or classify DNA sequences. Deep Learning has been efficiently used for classification of DNA sequences, using one-hot label encoding and Convolution Neural Networks (CNN)^22,23^, albeit the examples in literature are featuring DNA sequences of length up to 500 bps, only.

In particular, for the case of viruses, NGS genomic samples might not be identified by BLAST, as there are no reference sequences valid for all genomes, as viruses have high mutation frequency^24^. Alternative solutions based on deep learning have been proposed to classify viruses, by dividing sequences into pieces of fixed length, ranging from 300 bps^24^ to 3,000 bps^25^. However, this approach has the negative effect of potentially ignoring part of the information contained in the input sequence, that is disregarded if it cannot completely fill a piece of fixed size. The global impact of SARS-CoV-2 prompted researchers to apply effective alignment-free methods to the classification of the virus: For example, in^26^ the authors propose the use of Machine Learning Digital Signal Processing for separating the virus from similar strains, with remarkable accuracy. Nevertheless, there is no human-readable information that can be extracted from their black-box procedure, so the biological insight provided by their approach is limited.

Given the impact of the world-wide outbreak, international efforts have been made to simplify the access to viral genomic data and metadata through international repositories, such as the National Genomics Data Center (NGDC) repository^11^, the National Center for Biotechnology Information (NCBI) repository^27^ and the Global Initiative on Sharing All Influenza Data (GISAID) repository^28^, expecting that the easiness to acquire information would make it possible to develop medical countermeasures to control the disease worldwide, as it happened in similar cases earlier^29–31^. Thus, taking advantage of the available information of international resources without any political and/or economic borders, we propose an innovative system based on viral gene sequencing.

Using a CNN to separate Coronaviruses belonging to different strains^32^, including SARS-CoV-2, we apply techniques inspired by explainable AI in computer vision to discover representative cDNA sequences that the network uses to classify SARS-CoV-2. We then validate the discovered sequences on datasets not used during the training of the CNN, and show how to exploit them to create a novel, highly informative set of sequence features (e.g. viral sequences). Such sequences can be later inspected and analyzed by human experts. Experimental results show that the new set of sequence features leads traditional, simple classifiers, to correctly assess SARS-CoV-2 with remarkable accuracy (> 99%). A few of the discovered sequences also possess the correct characteristics for potentially becoming primers, as just checking for their presence in samples is enough to specifically identify SARS-CoV-2 (Fig. 1). Laboratory testing on the most promising sequences identified showed that the primers found by our approach can be a viable alternative to the commonly adopted primers at the time of writing.

**Figure 1.**
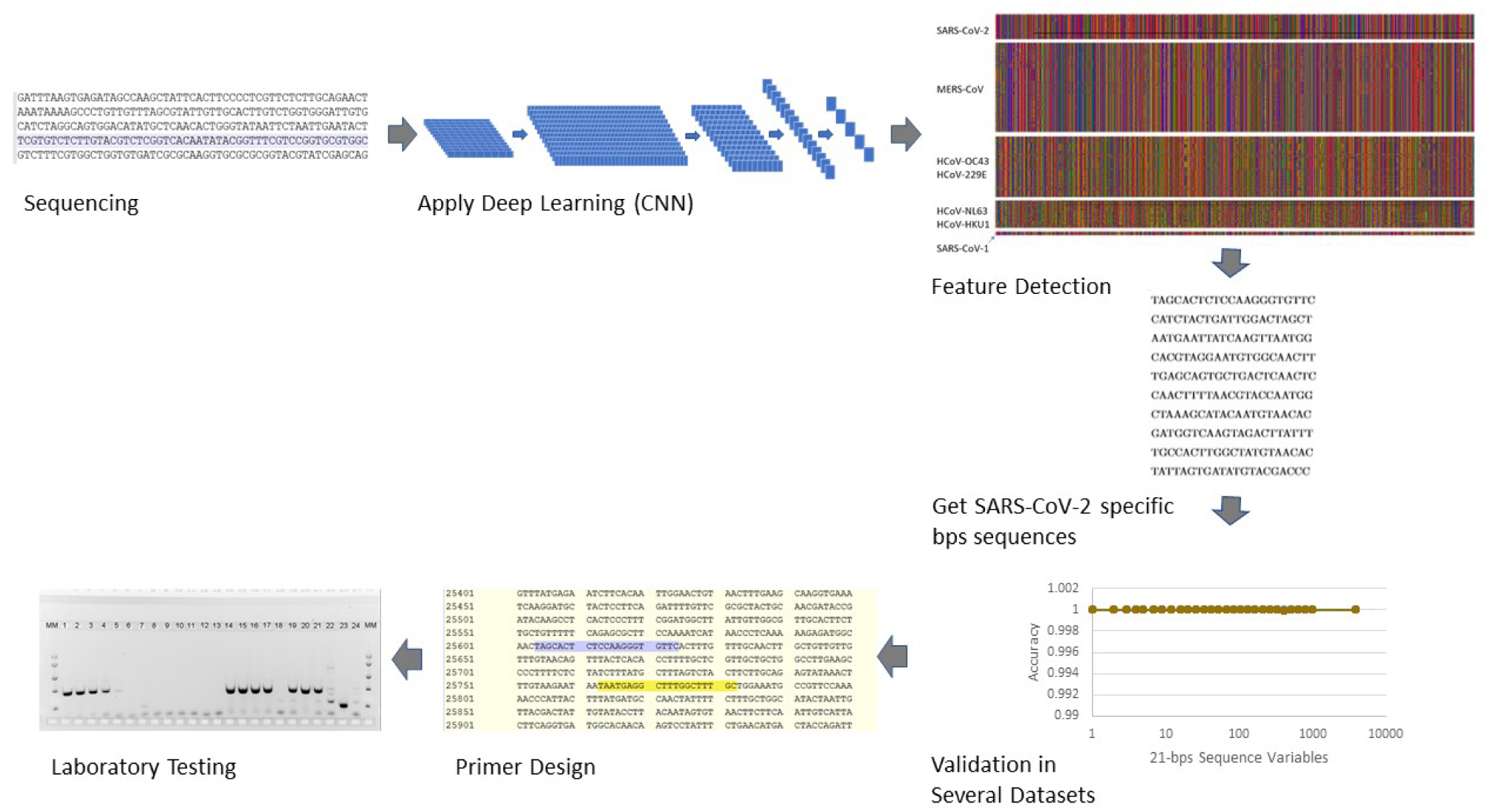
Overall procedure to find the specific SARS-CoV-2 21-bps RNA sequences to create a primer set.

## Results

### Identifying SARS-CoV-2

Summarizing the results of experiments 1-4 (Fig. 3), we discovered 12 meaningful 21-bps sequences that best characterize SARS-CoV-2. For all the analyzed data, these sequences appear only in SARS-CoV-2 samples and not in any other viruses, as summarized in Table 1. Remarkably, our results outperform earlier publications using machine learning for identifying SARS-CoV-2 (see for example^26^), with the added benefit of producing human-readable results instead of a plain black box classifier.

**Table 1.**
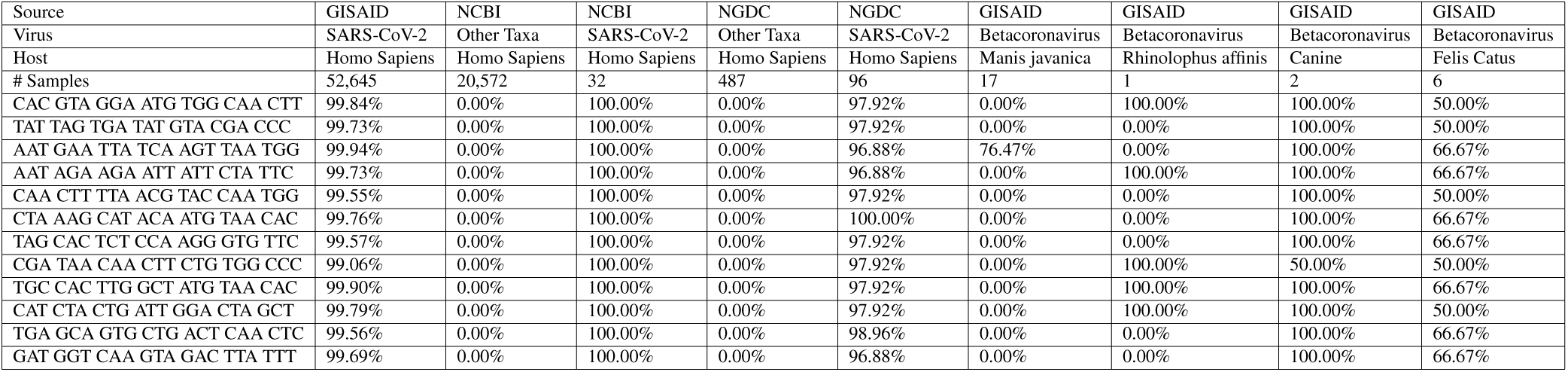
Percentage of appearance for each of the 12 discovered 21-bps sequences across the different datasets, and comparison to similar viruses in nature and other hosts.

### Laboratory validation of the candidate primer set

We calculated the frequency of appearance of different primer sets’ sequences used in SARS-CoV-2 RT-PCR tests developed by WHO referral laboratories and compared it to our primer design in the dataset from the GISAID (Table 2) repository. All of the sequences have a frequency of appearance of > 99%, with the exception of CHINA-CDC-N-F with a 68.52%. This is consistent with the percentage of genomes with mutation in the primer region in the GISAID latest update summary of August 11^*th*28^. In the analysis of specificity in silico, we compared all the primers sets’ sequences with the NCBI-B and NGDC dataset, the results show that HKU-N-F, HKU-N-R, Charite-E-F, Charite-E-R and US-CDC-N2-F are not specific to SARS-CoV-2 as they detect SARS-CoV-1 too. The rest of the sequences, including our design, only appear in SARS-CoV-2. Thus, in summary from 8 different primer sets, 3 of them are not specific to SARS-CoV-2, and from the remaining 5, considering frequency of appearance only, our design is in 3rd best option calculated with the lower limit.

**Table 2.**
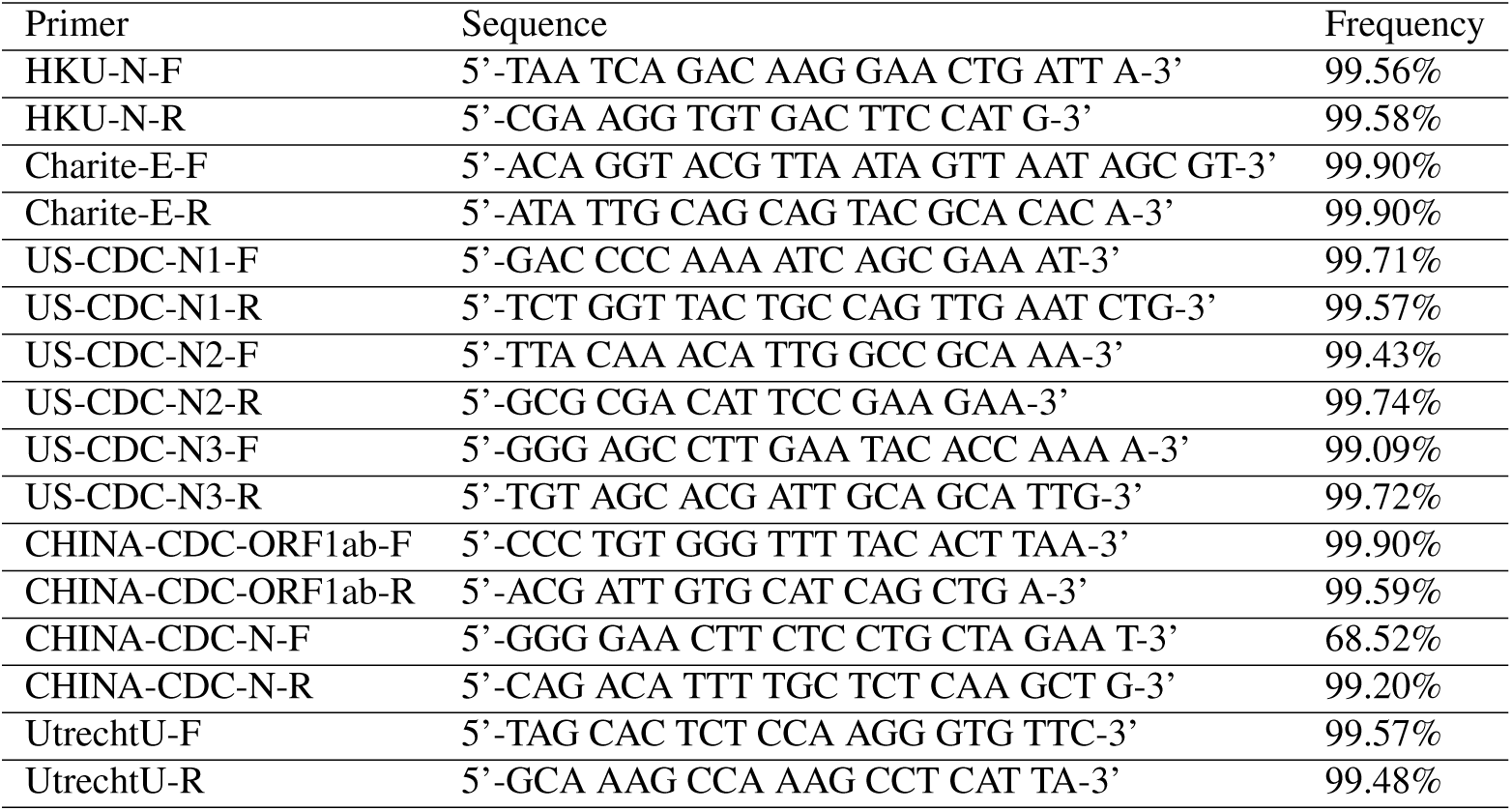
Frequency comparison for different sequences in primer sets suggested at the GISAID repository. Here, we denominated our design as UtrechtU.

To validate the data obtained in silico by laboratory methods a conventional PCR was performed on cDNA obtained from RNA from SARS-CoV-2 and other human coronaviruses. In addition, RNAs from nasopharyngeal swabs from six patients previously diagnosed with SARS-CoV-2 infection and four patients negative for SARS-CoV-2 by routine diagnostic method^5^ were analyzed with the same conventional PCR (Fig. 2). Different dilutions of SARS-CoV-2 RNA were detected with similar sensitivity compared to the diagnostic reference assay. (Fig. 2 lanes 1-8). Our candidate primer set exclusively detected SARS-CoV-2 and did not amplify RNA from other human coronaviruses (Figure 9, lanes 9-14). The candidate primer set was able to detect SARS-CoV-2 RNA from patient samples previously found positive for SARS-CoV-2, but not in patients previously found negative (Fig. 2, lanes 15-24). Although further validation will be required to develop this candidate primer set into a diagnostic assay, our results clearly demonstrate the power of our method to select potential sequences for further validation.

**Figure 2.**
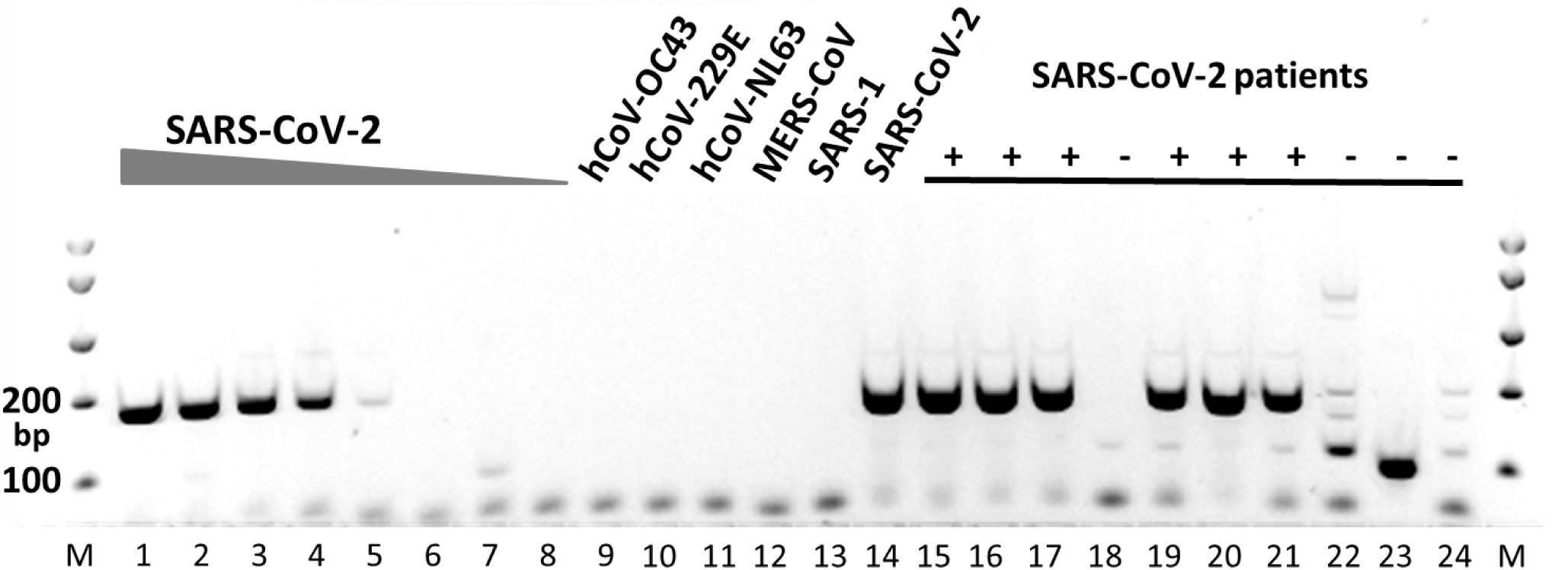
Laboratory validation of the candidate primer set by conventional PCR. MM, molecular weight marker; Lanes 1-8, 10-fold dilutions of SARS-CoV-2 RNA (corresponding to Ct values 26 to 39 in the diagnostic reference assay); Lanes 9-14, RNA from different human coronaviruses (hCoV-OC43, hCoV-229E, hCoV-NL63, MERS-CoV, SARS-1, SARS-CoV-2 respectively); Lanes 15, 16, 17, 19, 20, 21, patient samples previously found positive for SARS-CoV-2; Lanes 18, 22, 23, 24, patient samples previously found negative for SARS-CoV-2.

**Figure 3.**
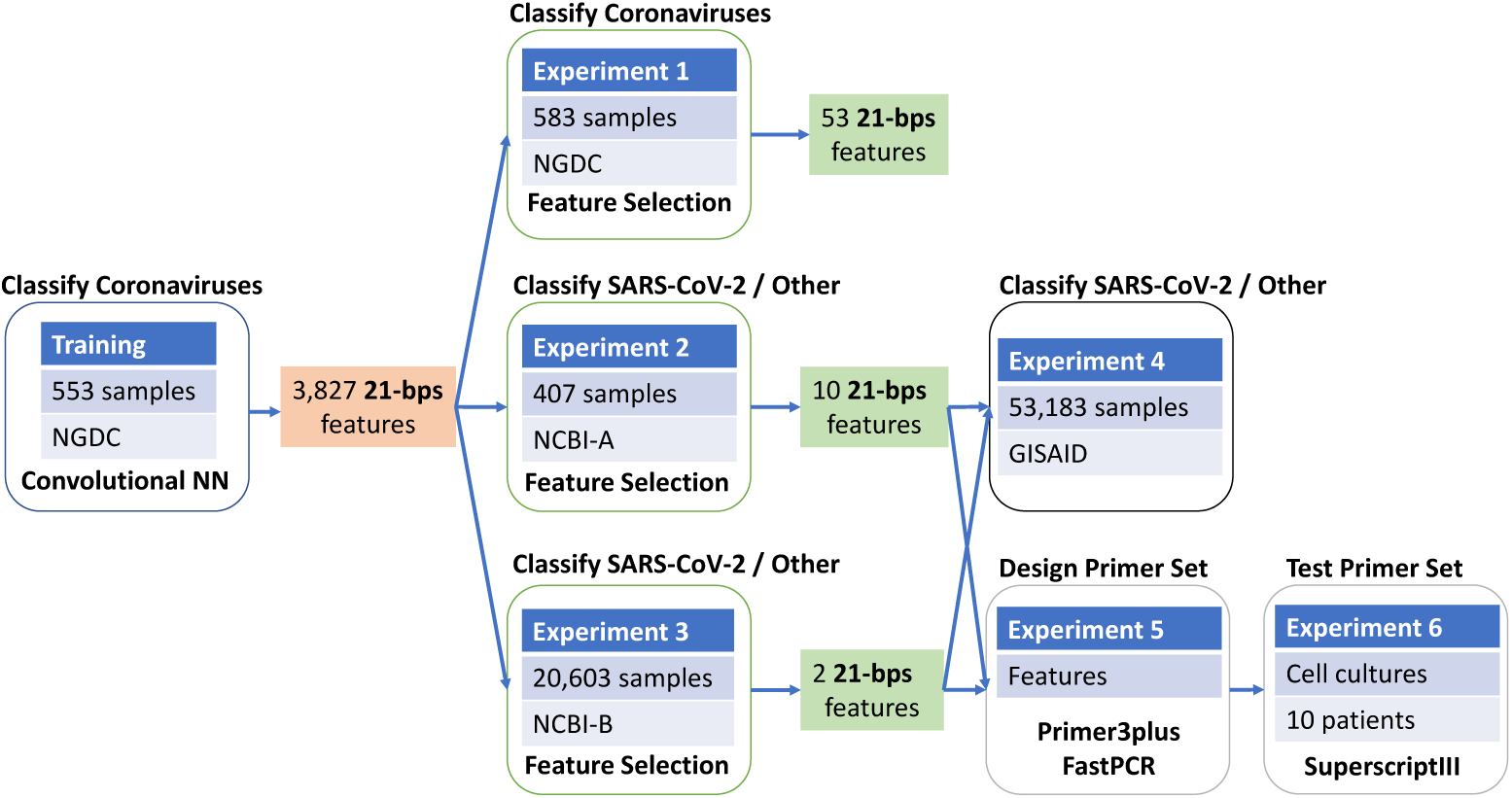
Summary of the different experiments, and corresponding datasets used.

## Discussion

Being able to reliably identify SARS-CoV-2 and distinguish it from other similar pathogens is important to contain its spread. The time of processing samples and the availability of reliable diagnostic tests is a challenge during an outbreak. Developing innovative diagnostic tools that target the genome to improve the identification of pathogens, can help reduce health costs and time to identify the infection, instead of using unsuitable treatments or testing. Moreover, it is necessary to perform an accurate classification to identify the different species of Coronavirus, the genetic variants that could appear in the future, and the co-infections with other pathogens.

Given the high transmissibility of the SARS-CoV-2, the proper diagnosis of the disease is urgent, to stop the virus from spreading further. Considering the false negatives given by the standard RT-qPCR detection, better implementations such as using deep learning are necessary in order to properly detect the virus. While the accuracy of current RT-qPCR testing is around 70%, and CT scans with deep learning go up at 83%, we believe that the use of the sequences detected by a CNN-based methodology has the potential to improve the accuracy of the diagnosis.

Our results, show that by targeting one out of the 12 selected 21-bps specific sequences, we are able to distinguish SARS-CoV-2, from any other virus (> 99%). Further testing is necessary to confirm these promising results so it is essential to create multidisciplinary groups that work to stop the outbreak. Finally, as an interesting remark, by comparing the discovered sequences against other hosts, we noticed that from the 12 sequences exclusive to SARS-CoV-2, one of them appears in 13 of 17 samples from *Manis Javanina*. In contrast, 5 of the sequences of SARS-CoV-2 appear in the only sample available from *Rhinolophus Affinis* and 11 out of 12 in 2 *Canine* samples(Table 1). This is consistent with the findings of Zhang et al.^33,34^, and could point to the zootonic origin of the virus. Nevertheless, more data is necessary.

As a result of the high density populations, and ever growing interaction between people, it is possible that other pandemics may occur. We believe that our methodology has a substantial added value over traditional methods, because it is a fast method and only limited set of viral sequencing data is needed. Moreover, this procedure led to a primer set with a very high specificity for SARS-CoV-2 with at least the same accuracy as the best primers sets in the world developed by WHO referral laboratories. Thus, thinking forward, our methodology can be applied in future viral pandemics to speed up the development of accurate detection methods for diagnosis and thereby contribute to limit the spread of a virus.

## Methods

The CNN used during all the experiments is composed of one convolutional layer with 12 different filters or weights (each with window size 21) with maxpooling (with pool size and stride 148), a fully connected layer (196 rectified linear units with dropout probability 0.5), and a final softmax layer with 5 units, to differentiate the different classes of Coronavirus strains. The optimized used is Adaptive Momentum (ADAM)^35^, with learning rate 10^−5^ and a batch size of 50 samples, run for 1,000 epochs^32^.

The convolutional layer of the network, in simple terms, is analyzing subsequences of 21 base pairs that can appear in different points of the virus genome. We selected 21 as designed primers for RT-PCR tests have a length of 18-22 bps normally. The pool size of the maxpooling represents the interval in which a specific 21-bps sequence can be recognized (in this case, 148 positions). Through the training process, the convolutional layer is *de-facto* learning new features to characterize the problem, directly from the data. In this specific case, the new features are 21-bps sequences that can more easily separate different virus strains. By analyzing the result of each filter in a convolutional layer, and how its output interacts with the corresponding max pooling, it is possible to detect human-readable sequences of base pairs that might provide domain experts with relevant information. It is important to notice that these sequences are not bound to specific locations of the genome; thanks to its structure, the CNN is able to detect them and recognize their importance even if their position is displaced in different samples.

We downloaded 583 sequences (*.*fasta* files) from the NGDC on March 15^*th*^,2020 (Table 3). We left out 30 SARS-CoV-2 sequences and then, we divided the rest of the data into 80% training, 10% validation, 10% testing. The trained CNN described above obtained a mean accuracy of 98.73% in a 10-fold cross-validation. Once the network is trained, in a first analysis, we plot the inputs and outputs of the convolutional layer, to visually inspect for patterns. As an example, in Fig. 4a we report the visualization of the first 1,250 bps of each of the 553 samples from the NGDC^11^ repository.

**Table 3.** Organism, assigned label, and number of samples in the unique sequences for the NGDC repository (left) and query: *gene=“ORF1ab” AND host=“homo sapiens” AND “complete genome”* in the NCBI repository (right). We use the NCBI organism naming convention^36^.

| Organism | Label | Number of Samples | Organism | Label | Number of Samples |
| --- | --- | --- | --- | --- | --- |
| SARS-CoV-2 | 0 | 96 | SARS-CoV-2 | 0 | 68 |
| MERS-CoV | 1 | 240 | MERS-CoV | 1 | 180 |
| HCoV-OC43 | 2 | 132 | HCoV-OC43 | 1 | 105 |
| HCoV-229E | 2 | 22 | HCoV-NL63 | 1 | 29 |
| HCoV-EMC | 2 | 6 | HCoV-HKU1 | 1 | 13 |
| HCoV-4408 | 2 | 2 | HCoV-4408 | 1 | 2 |
| HCoV-NL63 | 3 | 58 | HCoV-229E | 1 | 3 |
| HCoV-HKU1 | 3 | 17 | HCoV-EMC | 1 | 3 |
| SARS-CoV | 4 | 7 | HAstV-VA1 | 1 | 1 |
| SARS-CoV P2 | 4 | 1 | HAstV-BF34 | 1 | 1 |
| SARS-CoV HKU-39849 | 4 | 1 | HMO-A | 1 | 1 |
| SARS-CoV GDH-BJH01 | 4 | 1 | HAstV-SG | 1 | 1 |
| Total Samples | - | 583 | Total Samples | - | 407 |

**Figure 4.**
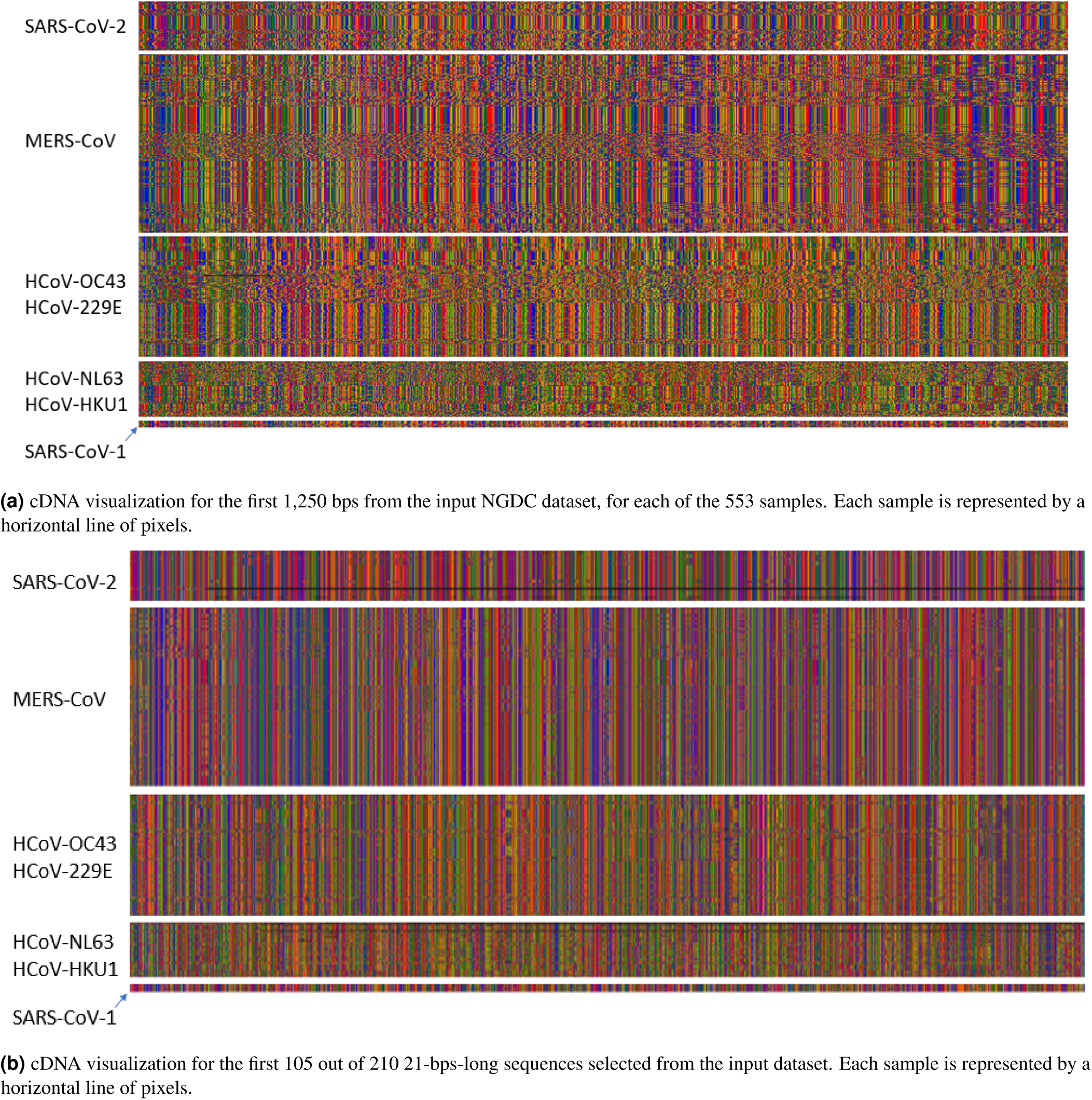
Input 4a, and output 4b of the methodology in colored pixels represent bases: G=green, C=blue, A=red, T=orange, missing=black. The data is separated by class (Table 3) SARS-CoV1: SARS-CoV, SARS-CoV P2, SARS-CoV HKU-39849 and SARS-CoV GDH-BJH01. For visualization purposes we do not show HCov-EMC and HCoV-4408, given the number of examples.From visual inspection we can see the similarity of the patterns between the classes.

Each filter slides a 21-bps window over the input, and for each step produces a single value. The output of a filter is thus a sequence of values in (0, 1). The output of the max pooling for each of the 12 filters is then further inspected for patterns. It is noticeable how samples belonging to different classes can be already visually distinguished. At this step, we identify filter 0 as the most promising, as it seems to focus on a few relevant points in the genome, that could correspond to meaningful cDNA sequences.

Given this data, it is now possible to identify the 21-bps sequences that obtained the highest output values in the max pooling layer of filter 0, in a section of 148 positions. This process results in 210 (31,029 divided by 148) *max pooling features*, each one identifying the 21-bps sequence that obtained the highest value from the convolutional filter, in a specific 148-position interval of the original genome: The first max pooling feature will cover positions 1-148, the second will cover position 149-296, and so on. We graph the whole set of max pooling features for the complete data 4,410 (210*21), Fig. 4b. The CNN architecture, and the visualization of the filter, and max pooling are available in the supplementary material section 1.

Analyzing the different sequence values appearing in the max pooling feature space, a total of 3,827 unique 21-bps cDNA sequences, that can potentially be very informative for identifying different virus strains. For example, sequence **AGG TAA CAA ACC AAC CAA CTT** is only found inside the class of SARS-CoV-2, in 59 out of 66 available samples. Sequence **CAC GAG TAA CTC GTC TAT CTT** is present again only in SARS-CoV-2, in 63 out of the 66 samples.

The combination of the convolutional and max pooling layer allows the CNN to identify sequences even if they are slightly displaced in the genome (by up to 148 positions). Thus, we create a table of feature appearance of each of the sequences selected from the previous step. This results, in just a set of feature to differentiate SARS-CoV-2 from other viruses.

The experiments presented in the following subsections to validate our method have different objectives and make use of different datasets. A summary of all the experiments and datasets used is shown in Figure 3.

### Identifying SARS-CoV-2

#### Experiment 1: Validation on the NGDC dataset

We downloaded the dataset from the NGDC repository^11^ on March 15^th^ 2020. We removed repeated sequences and applied the procedure to translate the data into the sequence feature space. This leaves us with a frequency table of 3,827 features (21-bps sequences) with 583 samples (Table 3 (left)). Next, we ran a state-of-the-art feature selection algorithm^37, 38^, to reduce the sequences needed to identify different virus strain to the bare minimum. Remarkably, we are then able to correctly differentiate all the coronavirus (MERS-CoV, SARS-CoV-2, SARS-CoV-1, etc) samples using only 53 of the original 3,827 sequences, obtaining a 100% accuracy in a 10-fold cross-validation with a simpler and more traditional classifier, such as Logistic Regression. The list of the 53 features is available in the supplementary material section 2.

#### Experiment 2: Validation on the NCBI dataset

We downloaded data from NCBI^27^ on March 15^th^ 2020, with the following query: *gene=“ORF1ab” AND host=“homo sapiens” AND “complete genome”*. The query resulted in 407 non-repeated sequences (Table 3 (right)). We call this dataset NCBI-A, where 68 sequences belong to SARS-CoV-2. Then, we applied the procedure to translate the data into the set of sequence features, and we run the same state-of-the-art feature selection algorithm^37^. The result is a list of 10 different sequences (Table 4), for which just checking for their presence is enough to differentiate between SARS-CoV-2 and other viruses in the dataset, with a 100% accuracy. Each of the sequences, in fact, only appears in SARS-CoV-2 samples.

**Table 4.**
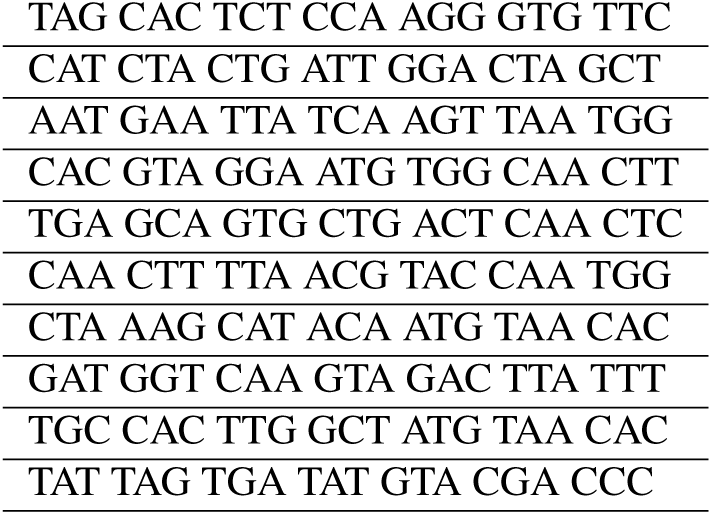
Sequences that only exist in SARS-CoV-2, that help differentiate between the virus and other taxa.

#### Experiment 3: Further validation on the NCBI dataset

We downloaded data from NCBI^27^ on March 17^th^ 2020, with the following query: *“virus” AND host=“homo sapiens” AND “complete genome”*, restricting the size from 1,000 to 35,000 bps (NCBI-B). The query returns 20,603 samples, of which only 32 belong to SARS-CoV-2, and 20,571 are from other taxa, including Hepatitis B, Dengue, Human immunodeficiency, Human orthopneumovirus, Enterovirus A, Hepacivirus C, Chikungunya virus, Zaire ebolavirus, Human respirovirus 3, Orthohepevirus A, Norovirus GII, Hepatitis delta virus, Mumps rubulavirus, Enterovirus D, Zika virus, Measles morbillivirus, Enterovirus C, Human T-cell leukemia virus type I, Yellow fever virus, Adeno-associated virus, rhinovirus (A, B and C), for a total of more than 900 viruses. Then, we applied the procedure to translate the data into the sequence feature space and run the feature reduction algorithm^37^. This results in 2 extra sequences of 21 bps: just by checking for their presence, we are able to separate SARS-CoV-2 from the rest of the samples with a 100% accuracy. The sequences are: **AAT AGA AGA ATT ATT CTA TTC** and **CGA TAA CAA CTT CTG TGG CCC**.

#### Experiment 4: Validation on the GISAID dataset

From the GISAID repository^28^, we downloaded 53,183 sequences available on August 10^th^, for SARS-CoV-2, from different countries, from there 52,645 have as < 1% Ns, high coverage and *host=“homo sapiens”*. Then, we calculated the frequency table of the 21-bps sequences obtained from experiments 2 and 3, to verify which sequences remain and could be used for detection. The appearance frequency of the target sequences among the samples in the GISAID dataset is reported in Table 1, second column. In addition, we downloaded 26 sequences from GISAID repository of other hosts (*manis javanica, rhinolophus affinis, canine* and *felis catus*) to make a comparison in the sequences from experiment 2 and 3.

### Laboratory validation of the candidate primer set

#### Experiment 5: Design of the candidate primer set

After the analysis carried out on the deep learning model, we ran an analysis with Primer3plus^39^, to see which of the sequences could be used as a forward primer, using sample NCBI NC045512.2 as the reference SARS-CoV-2 sequence. We uncover the sequence **TAG CAC TCT CCA AGG GTG TTC** that shows a frequency of appearance of 99.57% in viral genomes available from different countries in GISAID^28^ and 100.0% in the NCBI^27^ datasets. Using the reference SARS-CoV-2 sequence, we identify that this discovered sequence is located between nucleotides 25,604 and 25,624 in the ORF3a gene. In SARS-CoV, this gene encodes a protein of 274 aa, that is related with necrotic cell death^40^, chemokine production like interleukin 8 (IL-8) and RANTES/CCL5, NF*κB* activation resulting in an inflammatory response^41^ and may play an important role in the virus life cycle^42^. We design a specific primer set for detection of SARS-CoV-2 using Primer3plus^39^. We use **TAG CAC TCT CCA AGG GTG TTC** as forward primer and **GCA AAG CCA AAG CCT CAT TA** as reverse primer, obtaining an amplicon size of 179 bps. Then, we run an *in-silico PCR* test using FastPCR 6.7^43^ with default parameters in NC045512.2 used as a reference SARS-CoV-2 sequence, this yields *Tm* = 56.2°*C* for the forward primer, *Tm* = 53.1°*C* for the reverse primer and *Ta* = 58°*C*.

In addition, we calculated the frequency of appearance of different primers sets’ sequences used in SARS-CoV-2 RT-qPCR tests developed by WHO referral laboratories and compared it to our primer design sequences in 52,645 sequences from the GISAID repository and the 583 samples of different coronaviruses from the NGDC dataset from experiment 1. The used primers set are developed by University of Hong Kong (HKU-N); Charite, Berlin, Germany (Charite-E); US-CDC, United States (US-CDC-N1,US-CDC-N2,US-CDC-N3) and China CDC, China (China-CDC-ORF1ab, China-CDC-N) (Table 9). We selected this primers as they are the ones more commonly used as stated in the GISAID status update of August 11, 2020. We do not consider degenerate primer sets.

#### Experiment 6: Validation of the candidate primer set in biological samples

Viral RNA was isolated from cell-cultured SARS-CoV-2, SARS-1, MERS-CoV, hCoV-NL63, hCoV-OC43, hCoV-229E, and from nasopharyngeal swabs from *n* = 10 patients by MagNA Pure LC (Roche Diagnostics, The Netherlands) using the total nucleic acid isolation kit. The RNA was converted into cDNA using SuperscriptIII (Thermo-Fisher Scientific, USA) and random hexamers. Subsequently, conventional PCR was performed on the cDNA using HotStar Taq DNA polymerase (Qiagen, The Netherlands) with 400nM forward primer (5’-**AG CAC TCT CCA AGG GTG TTC**-3’) and 400nM reverse primer (5’-**GCA AAG CCA AAG CCT CAT TA**-3’) and the following cycling conditions: 15 min at 95°C, followed by 40 cycles of 1 min. at 95°C, 1 min. at 5 °CC and 1 min. at 72°C. The PCR products were visualized by electrophoresis. The same RNA was used in a diagnostics reference assay by Corman et al.^5^ and the Cycle threshold values form this reference assay were used for estimating sensitivity.

## Acknowledgements

The study was approved by the Medical Ethical Commission of the Erasmus MC (MEC-2015-306).

## Author contributions statement

LMM, CAP made the biological analysis, and primer design. ALR and AT made the programming, data collection and experiments in silico. DM and RM made the PCR validation. EC, ADK and JG made the experiment and study design. All the authors contributed to the writing.

## Notes

http://github.org/albertotonda/deep-learning-coronaviru

### Competing Interest Statement

The authors have declared no competing interest.

### Summary of Updates

Update Dataset Comparison to known Primers

