## Additional Information for "Classification and Specific Primer Design for Accurate Detection of SARS-CoV-2 Using Deep Learning"

### 1. CNN ARCHITECTURE AND VISUALIZATION

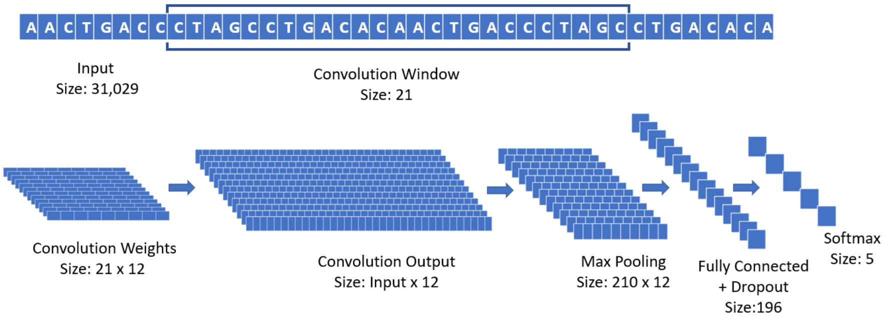


**Fig. S1.** CNN architecture to identify critical 21-bps sequences.

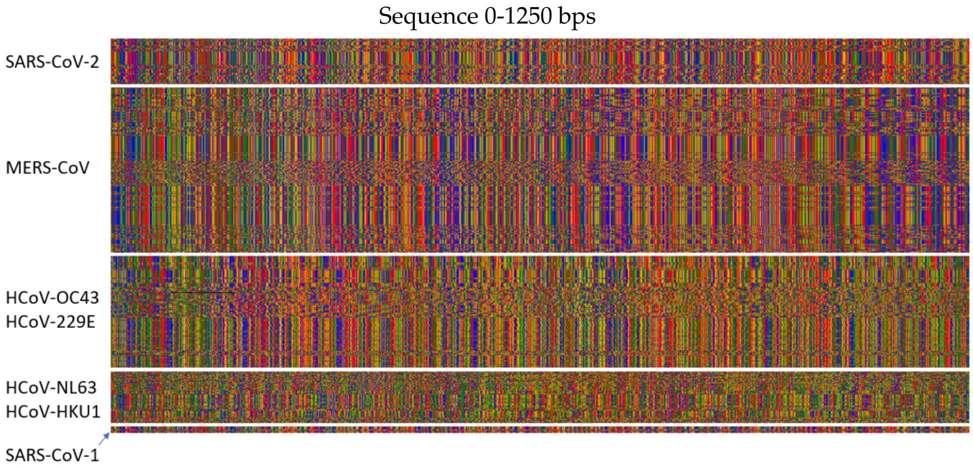


**Fig. S2.** cDNA visualization for the first 1,250 bps from the input NGDC dataset, for each of the 553 samples. Each sample is represented by a horizontal line of pixels. Colored pixels represent bases: G=green, C=blue, A=red, T=orange, missing=black. The data is separated by class SARS-CoV1: SARS-CoV, SARS-CoV P2, SARS-CoV HKU-39849 and SARS-CoV GDH-BJH01. For visualization purposes we do not show HCov-EMC and HCoV-4408, given the number of examples.

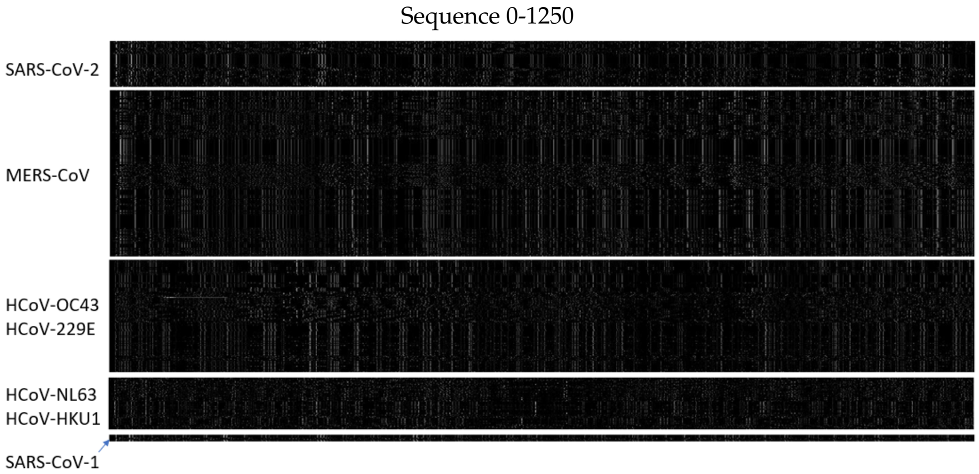


**Fig. S3.** The output of convolutional filter 0, for the input given in Fig. S2. The output of the filters is a series of continuous values in $(0, 1)$, here represented in grayscale, with higher values closer to white.

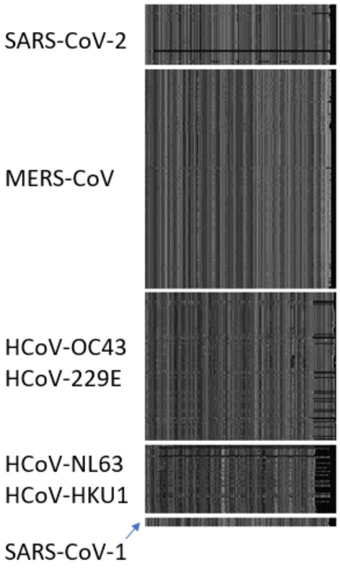


**Fig. S4.** Visualization of the output of the max pooling for the first filter of the CNN, with the data visualized in Fig. S3 in input. Different patterns for samples from different classes are recognizable from a simple visual inspection. Here samples from the same class appear grouped, one after the other, for clarity.

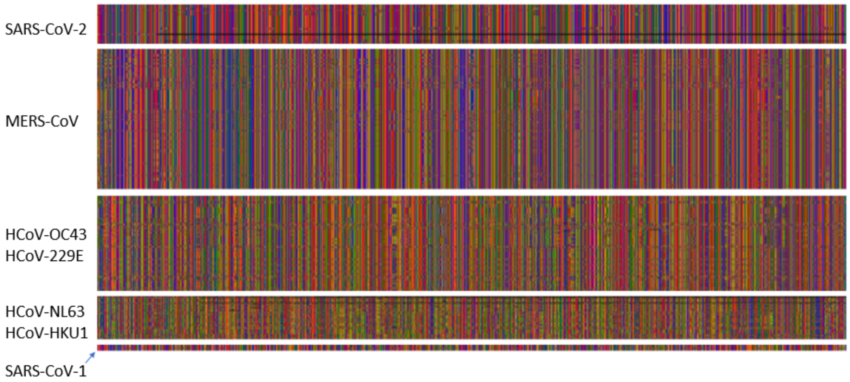


**(a)** 0 - 2,205 bps

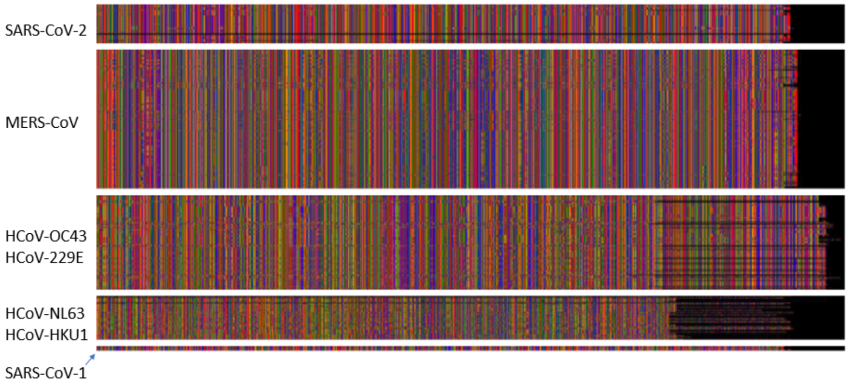


**(b)** 2,205 - 4,410 bps

**Fig. S5.** cDNA visualization for the selected 210 21-bps-long sequences selected from the input dataset. Each sample is represented by a horizontal line of pixels. Colored pixels represent bases: G=green, C=blue, A=red, T=orange, missing=black. We divide the whole information, for visualization purposes; from visual inspection we can see the similarity of the patterns between the classes.

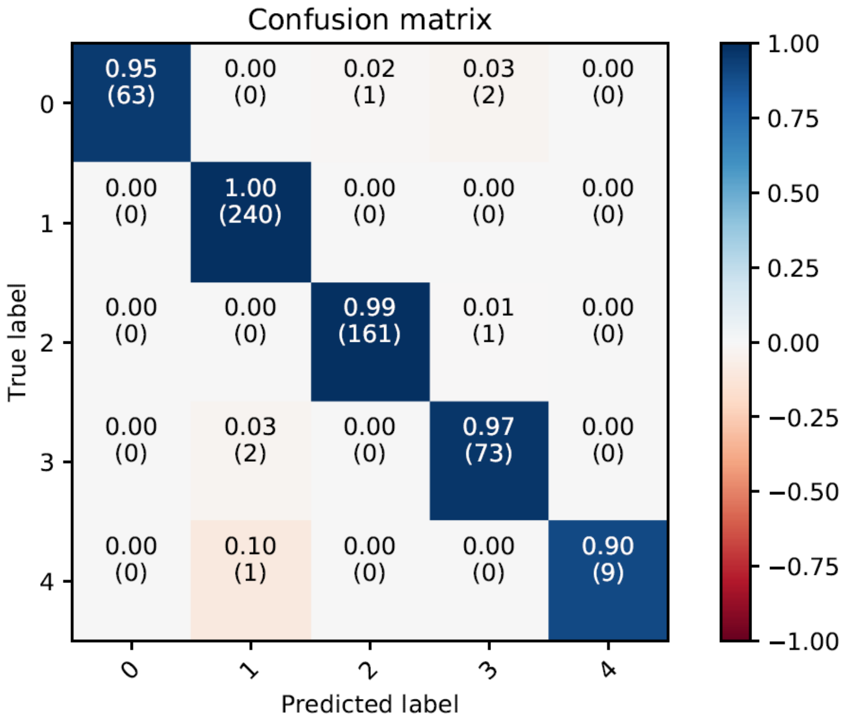


**Fig. S6.** Confusion Matrix Result of the CNN.

### 2. FEATURE LIST TO CLASSIFY DIFFERENT CORONAVIRUSES

**Table S1**

| | |
|---|---|
| TACAACATACCCTACTAATGT | CACCACTATTTGGGTTAACCC |
| TAGCATTAACCATGATTTGTT | GTTGAATGATCTTTGCTTCTC |
| GGTGATAAATTAGATCAGTTC | AGCGATAGGTTTTATCGACTT |
| CTAAAGCATACAATGTAACAC | CAAAAGTAACTAGTGCTATGC |
| TACCAGTGATTTAGGTAGTCT | TGCTTTAGATCAAGCTATTTC |
| AGGTCAAGAGCAAACTAACTG | TAACACTTACCTGGAGGGTTC |
| GAAGAGCTACCAGACGAATTC | GATTCTCAAATTTCTCATTAC |
| TGAATATAAATGATTGCACTC | CGGCTTGATTCAATTCAAGCC |
| GAAGGTAGTCGTGTGCCACAC | TATTAACAAGGACACTTATGC |
| TGTGGAGTATCCCATCATTTC | TGGGCTCTTGCCAACCAATTC |
| CTTCGGGAACGTGGTTGACCT | TATTGACTATAGACACTATTC |
| TGACTGCTAACAATCTAACTG | ATGCTATGGGCAAACCTGTGC |
| CTTGGTTCACCGCTCTCACTC | TATTGCTGACCTTGCTTGTGC |
| CAAAATCAGCGAAATGCACCC | TGACAGCTAACAATCTAACTG |
| GATTAATGATATGGTTTATTC | CAAAGTCTACTATGGTAATGC |
| TGTGTTTAGTCAAGTTGATTT | CAATGTTTATTGTTATAACAC |
| TGCTAACAATCTAACTGCACC | AAGAGAGAAGTTGCTTCATTT |
| GCTGAATAAGCATATTGACGC | TAAAGTTCATTTTTATTACCT |
| GGGTTGCAACTGAGGGAGCCT | TGGCTGCAACGTAACTGACGT |
| AATTGGAATGTGGAGTATCCC | CTGTGGCTACCTTGACGGCGC |
| CTTGACAGATTGAACCAGCTT | CAAATAGCACCTGTTCCAGCT |
| AGGCGGCAGTCAAGCCTCTTC | AATGTCTAAACTAAACGATGT |
| TATTTGCAATTCAGCTGTTGC | TGCTGATGATTCAGGTACTCC |
| GGGCCAGAAGCTGGACTTCCC | GACCTTAAATTCAGACAACGT |
| AGACGTGGTCCAGAACAAACC | CATGCTGTACCGACTCTCTTT |
| TATACTCAACTGTGTCAATAC | TACAGAATTTTAAGGAAACGC |
| TGGCCGCAAATTGCACAATTT | |
